# Representation learning applications in biological sequence analysis

**DOI:** 10.1101/2021.02.26.433129

**Authors:** Hitoshi Iuchi, Taro Matsutani, Keisuke Yamada, Natsuki Iwano, Shunsuke Sumi, Shion Hosoda, Shitao Zhao, Tsukasa Fukunaga, Michiaki Hamada

## Abstract

Remarkable advances in high-throughput sequencing have resulted in rapid data accumulation, and analyzing biological (DNA/RNA/protein) sequences to discover new insights in biology has become more critical and challenging. To tackle this issue, the application of natural language processing (NLP) to biological sequence analysis has received increased attention, because biological sequences are regarded as sentences and k-mers in these sequences as words. Embedding is an essential step in NLP, which converts words into vectors. This transformation is called representation learning and can be applied to biological sequences. Vectorized biological sequences can be used for function and structure estimation, or as inputs for other probabilistic models. Given the importance and growing trend in the application of representation learning in biology, here, we review the existing knowledge in representation learning for biological sequence analysis.

## 1. Introduction

Considerable advances in high-throughput sequencing have resulted in rapid data accumulation [1]. Although these modern technologies produce a large amount of data, they do not provide any interpretation or biological information. Thus, analyzing biological sequences, such as DNA/RNA/protein sequences, to make biological discoveries has become more critical and challenging. To tackle this issue, the application of natural language processing (NLP) to sequence analysis has attracted great attention in terms of biological sequences as sentences and k-mers in these sequences as words [2, 3].

NLP aims to allow computers to understand the contents of natural language, including the context, and accurately extract information and insights [4]. Natural language is composed of characters, such as the alphabet, and the meaning is constructed using grammar and semantics. In the same way, biological sequences can be regarded as sentences with different letters, and biophysical and biochemical rules define properties, such as the function and structure [5]. Biological sequences are consistent with natural language where characters define their meaning, and the meaning depends on the neighboring sequence. For example, whether the word “bank” in a sentence refers to a financial institution or raised portion of seabed depends on the context. Similarly, whether a part of an RNA sequence forms a secondary structure depends on its neighboring sequences. Thus, there are similarities between natural language and biological sequences, and it would be natural to apply NLP to a more in-depth understanding of the function and structure encoded in the biological sequence.

*Representation learning* is an essential step in NLP and indicates automatic systems to explore the representation of raw data, such as words or characters [6]. In general, the representation is provided as a real-valued vector, called *distributed representation*. Successful representation learning is expected to convert words into vectors while preserving their semantic similarity. For example, the names of foods, like “sushi” and “pizza,” should be converted into similar vectors and the names of organisms, such as “frog,” should be assigned entirely different vectors (Figure 1). In biological sequences, *N*-methyl-D-aspartate receptor and *α*-amino-3-hydroxy-5-methyl-4-isoxazolepropionic acid receptor, which are ionotropic glutamate receptors, are expected to be converted into similar vectors, whereas GFP, a fluorescent protein, is expected to be converted into a completely different vector. Thus, representation learning indicates the transformation from words to vectors while preserving the similarities and differences between words.

**Figure 1:**
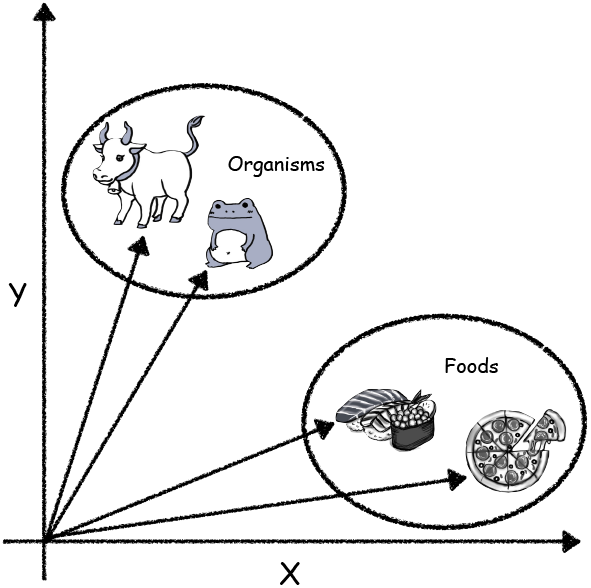
Ideal representation learning should convert the names of foods, such as “sushi” and “pizza,” into similar vectors and assign different vectors to the names of organisms, such as “cow” and “frog.”

Biological sequences vectorized by representation learning can be directly used for biological tasks, such as function and structure prediction. If the vector similarity between proteins is high, it can be inferred that they have similar functions and structures. Note that vector similarity/distance can be calculated using linear algebra operations, such as dot product, Euclidian distance, and cosine similarity. In particular, the successful encoding of words via representation learning has been recognized as an essential research area because the performance of NLP and deep learning depends on the quality of the representation [6]. Thus, a *good* representation of a biological sequence is critical for clustering, function, structure, and disorder prediction [2].

Given the significance and growing trend in the application of representation learning in biology (Figure 2), here, we describe a review of representation learning for sequence analysis. It should be noted that this review covers the application of representation learning to biological sequence analysis, and its use in biological literature and medical records is beyond the scope of this review. This review is organized as follows: Section 2 introduces the basic representation techniques for NLP. Section 3 provides a comprehensive survey of representation learning approaches for sequence analysis. Section 4 presents a summary and an outlook of representation learning applications in biological sequence analysis.

**Figure 2:**
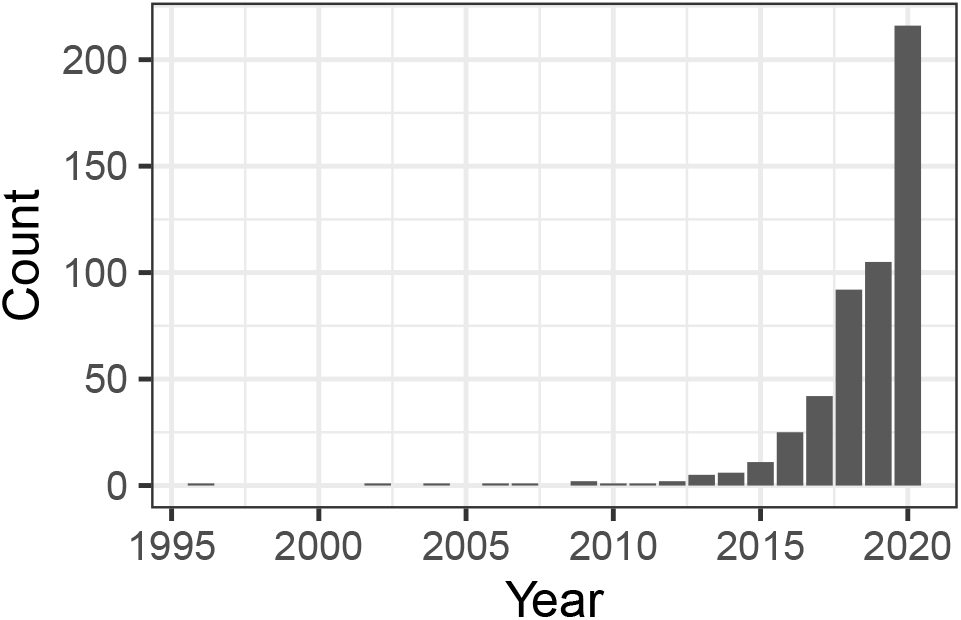
Change in the number of hits for the search term “representation learning” (with double quotation) in PubMed (https://pubmed.ncbi.nlm.nih.gov/).

## 2. Representation learning techniques

Currently, the acquisition of distributed representations of biological sequences is mainly achieved using neural networks developed in NLP. In representation learning for NLP, it is assumed that the words that appear in the same context should have similar meanings according to the *distribution hypothesis* [7]. Representation learning methods based on the distribution hypothesis attempt to vectorize words or phrases by training the neural networks with architectures specialized for capturing the relationships among words from a corpus, a set of documents. Various representation learning methods presented in this review are taken from neural-network-based language models specialized for biological sequences; thus, it is essential to understand the underlying architecture of the neural networks developed for NLP. In this section, we briefly summarize the development of basic representation learning techniques.

word2vec is the first successful method to obtain distributed representations using a neural network [8, 9]. There are two types of neural networks used in word2vec: skipgram model that predicts the words around the input word and a continuous bag-of-words model that predicts the target word from the surrounding words. Until the advent of word2vec, researchers used neural networks to describe the syntactic structure [10, 11]. The skip-gram model proposed by Mikolov attracted attention for its ability to capture not only grammatical correctness but also semantic features, as described in the introduction. word2vec with the skip-gram model acquires a distributed representation for each word by training the three-layer neural network, as shown in Figure 3. Given a sentence with *T* words and the *t*-th word *w_t_*, the model predicts the words present in the vicinity of *w_t_* in that sentence. The parameters to be estimated in the skipgram model include the weight matrix *X* to predict the *d* - dimensional hidden layer 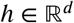 from the one-hot encoded input layer and weight matrix *Y* to predict the output from *h*. They are predicted using the formula described below:

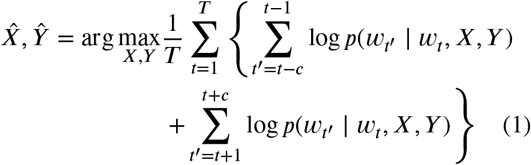

where *c* is a constant indicating the number of words farther away from *w_t_* that should be included in the prediction. The model performs the same operation for all sentences to complete training. In this case, the weight matrix *X* is a *V* × *d* matrix, where *V* is the number of words in the vocabulary. If *w_t_* is the *v*-th word in the vocabulary, we can obtain the distributed representation of the word *w_t_* as the *v*-th vector of the predicted *X* (i.e., 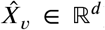). The word2vec representation has additive compositionality and has become famous for allowing intuitive operations, such as 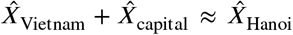, as shown previously [9]. Hence, word2vec succeeded in obtaining highly interpretable distributed representations for the first time and directed subsequent development in representation learning.

**Figure 3:**
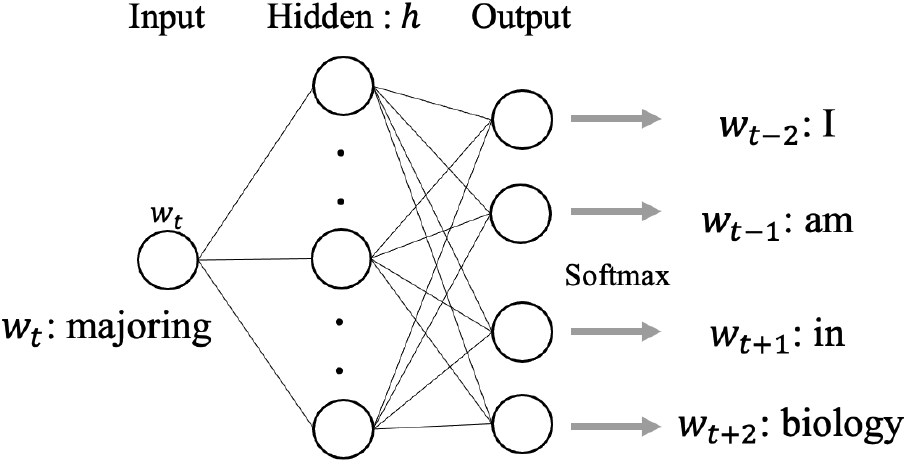
Skip-gram model used in word2vec. This neural network model has three fully connected layers: the input, hidden, and output layers. In this case, it attempts to learn the features from the sentence, “I am majoring in biology,” and predict the words surrounding *w_t_*, “majoring. “

The fact that word2vec captured semantic features was a remarkable breakthrough in representation learning, and various extended models based on word2vec were proposed. GloVe uses word co-occurrence matrices, which have been used in classical latent semantic analysis, such as singular value decomposition [12]. It shows higher semantic accuracy than word2vec. FastText is one of the embedding methods based on the skip-gram model [13]. It can train the model very quickly while maintaining the same accuracy as conventional methods. In addition, several methods were developed to obtain a distributed representation for each sentence (not word) based on the word2vec concept. doc2vec utilizes the paragraph vectors, which captures the context for each paragraph and provides the features for each sentence [14].

Although word2vec has enabled considerable progress in representation learning, it cannot express the semantic polysemy of words because it yields a single *d*-dimensional vector for a single lexicon, as mentioned above. For example, “right” that appears in “right to vote” and “turn right” differ in meaning, but they are embedded at the same point using word2vec. The approach to solving this problem is called word sense disambiguation in NLP [15], and it calls for architecture to consider the context and meaning of a sentence. In biological sequences, the context of a word in a sentence is equivalent to the role of a particular nucleic/amino acid in the whole sequence. Hence, the polysemy in biological sequences is critical, similar to that in natural languages. Here, we introduce two methods that can take such contexts into account: one that can achieve this by making the neural network recursive using a recurrent neural network (RNN) or long short-term memory (LSTM) [16] and another that uses the *attention* mechanism.

Embeddings from language models (ELMo) dissolves the polysemy problem using the model stacked with multiple bidirectional-LSTM (bi-LSTM) and yields the distributed representations by taking the linear weighted-sum of outputs of their hidden layers [17]. RNN and LSTM have been utilized mainly for sequential tasks, such as document generation and machine translation [18, 19]. In a language model with a simple forward LSTM, the occurrence probability of the *t*-th word in a sentence, *w_t_*, depends on the set of words that appear before *w_t_* (denoted as ***w***_1:*t*-1_). The model trains the parameters to maximize the joint probability for all words, {*w*_1_, …,*w_t_*, …,*w_T_*}. To calculate *p*(*w_t_*|**w**_1:*t*-1_), LSTM uses the hidden layer of *w_t_* (its output is denoted by *h_forward,t_* 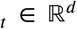), which depends on *w*_*t*-1_ and *h_forward,t-1_*. As the hidden layer is computed recursively depending on the word order, LSTM-based models allow context-aware learning. Previously [17], bi-LSTM modules were used in their language model called the bidirectional language model (bi-LM). bi-LSTM considers not only the forward but also backward word dependency. In a backward LSTM, the hidden layer of *w_t_* and its output *h_backward,t_* depend on *w_t+1_* and *h_backward,t+1_* as opposed to the case in forward LSTM. By considering word dependency in the backward direction, bi-LM can incorporate relationships among words that cannot be captured by the forward LSTM alone. bi-LM contains a stack of *L* bi-LSTM modules (see Figure 4), and all the modules are trained to maximize the joint probability of generating the entire sentence as follows:

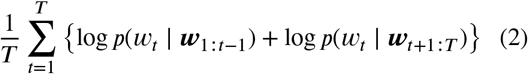

**Figure 4:**
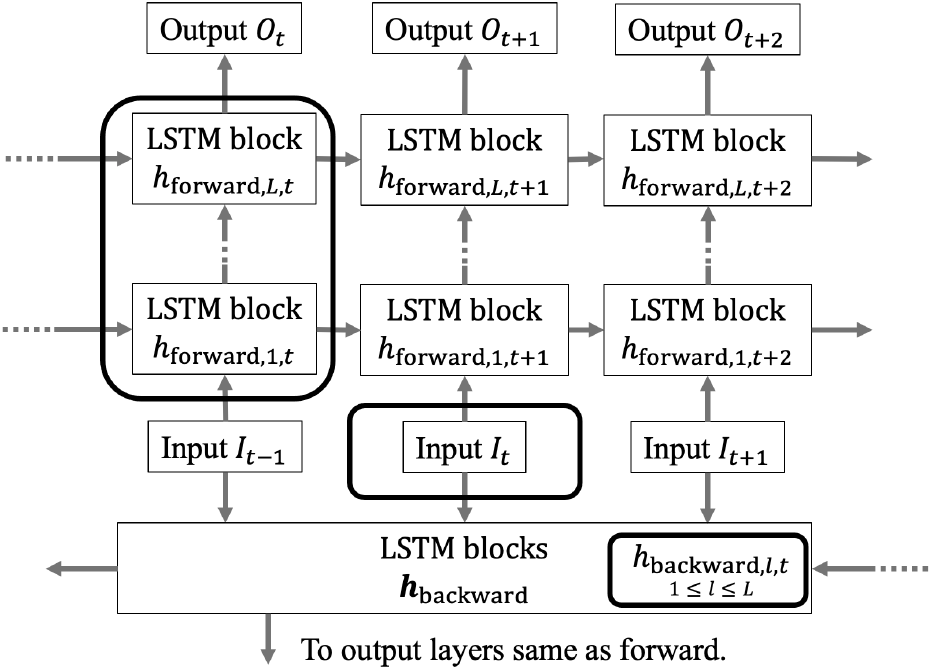
The graphical representation of a bidirectional language model. Input *I_t_* shows the embedding of the *t*-th word in the sentence, *w_t_*. The output, *O_t_*, is transformed to a probability with a softmax function, and all the modules are trained to maximize the observed probability of *w_t_*. 2*L* + 1 layers circled in squares with rounded corners are used to calculate ELMo_*t*_, the distributed representation for *w_t_*.

Finally, the distributed representation, ELMo, is obtained by taking the weighted-sum of outputs from 2*L* +1 layers, which are hidden layers for each LSTM module and an input embedding layer. ELMo succeeded in placing the same lexicon on different points in a high-dimensional space, depending on the context.

Another approach to solving the polysemy problem is to use the attention mechanism. In brief, attention quantifies the degree of correspondence between words [20, 21].

Neural networks with attention mechanisms have an attention weight that is obtained by calculating the association of hidden layers (e.g., using the inner product) for arbitrary combinations of words in sentences. If the two words used to compute the attention weight come from different sentences, this attention is called the source-target-attention; if they are from an identical sentence, it is called self-attention. Models that use attention weights in the forward propagation are highly expressive, and we can naturally introduce an attention mechanism to representation learning. Transformer, which implements attention mechanism and positional encoding [22] in Key-Value Memory neural network [23, 24] without conventional context-aware architectures, such as RNN or LSTM, has achieved the state-of-the-art accuracy in several tasks, including machine translation [25].

Bidirectional encoder representations from transformers (BERT) have multiple transformers with self-attention connected in series (see Figure 5) [26]. In the pre-training of BERT, the input is a set of tokens connecting two sentences. At this time, a part of the input words is masked, and the model predicts the masked words from the extracted features considering the context. In addition, the model performs a binary classification of whether the two input sentences are semantically consecutive. Similar to other methods, we can use the outputs of pre-trained transformer layers as the distributed representations of input sentences.

**Figure 5:**
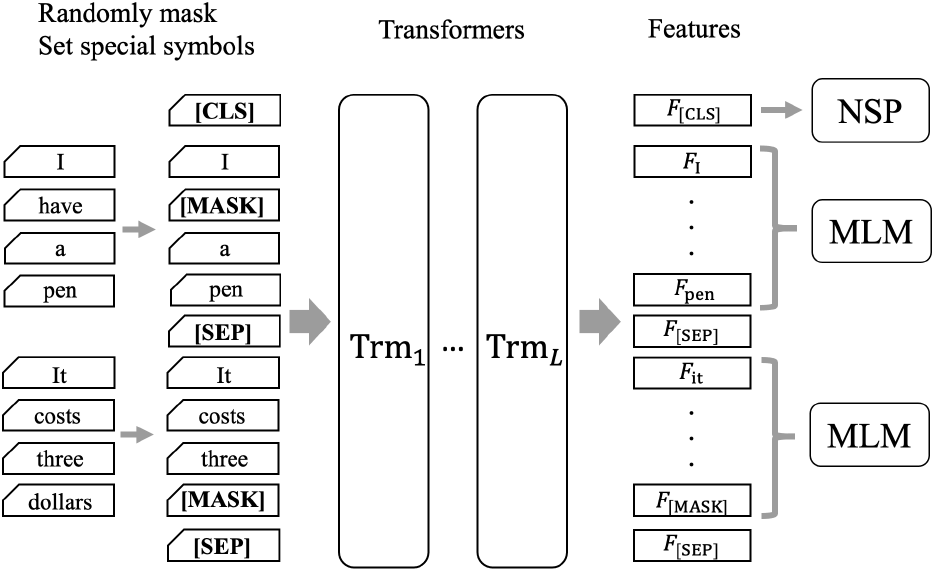
The graphical representation of Bidirectional encoder representations from transformers (BERT) architecture. Preparation of special tokens enables the model to extract features based on the self-attention of the whole sentence. BERT is trained with two tasks: masked language model (MLM) and next sentence prediction (NSP).

Neural networks with attention mechanisms, such as transformer and BERT, capture distal word associations better than conventional recursive models represented by RNN and LSTM [20, 27]. This is because, in recursive models, a hidden layer of a certain word depends on the hidden layers of the neighboring words only, and the contribution of distal words becomes small or converges to zero. In contrast, the use of attention is robust against such weight loss because the model always refers to the association of it with all words. This feature of BERT is attractive from the viewpoint of biology since distal interactions are important for structural prediction and other purposes. Another advantage of BERT is task-independent versatility. For instance, when we use ELMo, we should prepare a task-specific model to transfer the obtained distributed representations to other tasks. In contrast, with BERT, we can utilize the same architecture used in pre-training (as shown in Figure 5) without modification. Fine-tuning, which uses pre-trained hidden layers for initialization and optimizes the parameters for each task, has achieved state-of-the-art accuracy in many NLP tasks [26].

The main advantage of obtaining features through unsupervised learning is that it can retain versatility for transfer learning to various tasks. However, to build a specialized model for a specific task, representation learning in a *supervised* manner is also useful. `StarSpace` is a supervised learning method [28], which uses labeled documents as the training dataset, and embeds words and labels in the same space so that a label is close to words associated with it. Embedding with StarSpace allows for text classification, that is, prediction of labels used in the course of learning with higher accuracy than the other unsupervised methods, and provides highly interpretable vectors. As this example shows, supervised representation learning is also a practical option if the correct labels are known.

Since the development of word2vec in 2013, the field of representation learning in NLP has been growing at an astonishing pace. Considering the models based on transformer or BERT, several modern improved methods have continued to provide increased accuracy [29, 30]. Furthermore, similar to the big impact of the attention mechanism, the emergence of new concepts may also reconstruct the current paradigm of language modeling. These substantial developments in machine learning will be useful for bioinformatics and sequence analyses. As numerous examples are introduced in later sections, we believe that applying the latest representation learning techniques to biological sequences will lead to a discovery or elucidation of novel information in this domain.

## 3. Survey of representation learning applications in sequence analysis

We conducted an exhaustive survey, as shown in Table 1, for articles that met the following criteria: (i) Peer-reviewed and published in PubMed, except for BERT, which was recently published with a limited number of peer-reviewed articles; (ii) explicitly used a language model, such as word2vec or BERT; (iii) provided the source code or the model for repeatability or verification.

**Table 1.**
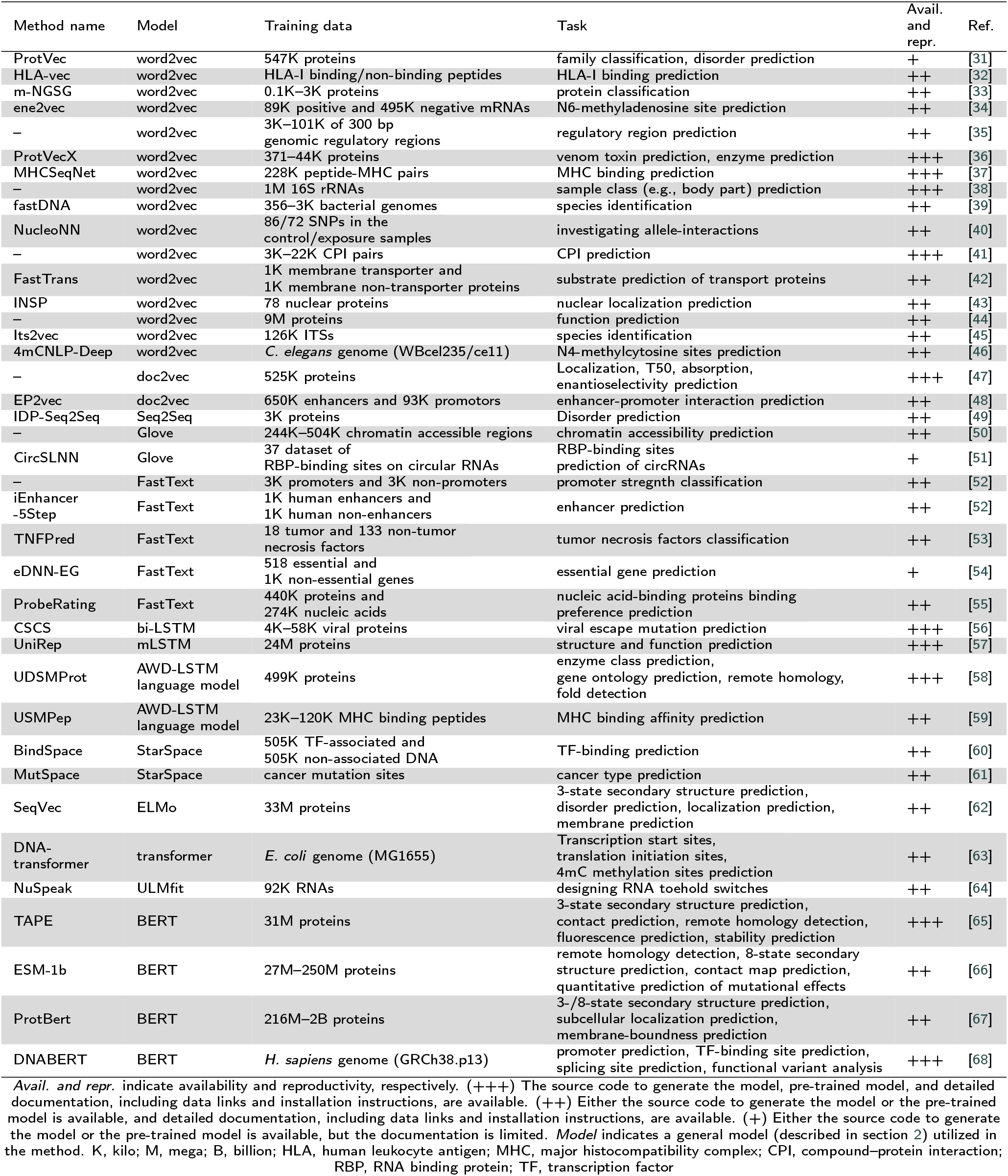
Comprehensive survey of representation learning application in biological sequences

### 3.1. Applications for structure/function prediction

**ProtVec** is the first model to use the embedding method for biological sequences [31]. This method regarded *3-mers* of amino acids as words and used 546,790 protein sequences from the Swiss-Prot database as the training dataset. Subsequently, word2vec using the skip-gram model was applied to the dataset, and 100-dimensional protein vectors were calculated. Originally, ProtVec was evaluated based on protein family classification and disordered protein prediction accuracies and it achieved high performance in both. Currently, ProtVec has also been utilized for predicting kinase activity [69] and gene function [70]. As ProtVec is a straightforward model, various extensions have been proposed. One of the extensions is seq2vec, which embeds not the k-mers of amino acids but the whole protein sequences [71]. Seq2vec utilizes doc2vec [14], an NLP method that embeds documents instead of words, which showed a higher performance than ProtVec in terms of protein family classification performance. Another extension is dna2vec [72], which embeds variable-length k-mers rather than fixed-length DNA k-mers using word2vec. ProtVecX is a similar method that uses word2vec to embed variable-length amino acid k-mers [36].

**SeqVec** is the first model that uses ELMo to achieve amino acid representation based on the whole protein sequence [62]. ELMo was applied to the UniRef50 dataset, which contains 33M proteins with 9.6G residues, regarding single amino acids as words. The extracted sequence profile was then fed to the per-residue prediction and per-protein prediction. With and without the evolutionary information, the model accurately predicted the secondary structure, disorder, localization, and membrane binding. The performance did not exceed that of the state-of-the-art methods [73, 74]. However, it was better than ProtVec [31] which is a context-independent model. In some tasks, such as protein function prediction, it outperformed one-hot encoding of k-mer-based embeddings and showed the competitive results obtained using ELMo [75].

**UDSMProt** is another language model representation extractor using a variant of LSTM [58]. The structure used was called AWD-LSTM [76], which is a three-layered bi-LSTM that introduces different types of dropout methods to achieve accurate word-level language modeling. UDSMProt was initially applied to the Swiss-Prot database and then fine-tuned for specific tasks, such as enzyme commission classification, gene ontology prediction, and remote homology detection. UDSMProt showed that upon pre-training with external data, the model performed as well as the existing methods that were tailored to the task using a position-specific scoring matrix (PSSM) and outperformed them in two out of three tasks. In addition, it demonstrated that utilizing pre-training information can compensate for the lack of data, compared to the case where PSSM information is provided. These results and extensions, such as USMPep, which revealed the ability to successfully predict MHC class I binding [59], imply that language models can efficiently contextualize and achieve word-based representation.

**ESM-1b** is a BERT-based model trained on a massive biological corpus, particularly amino acid sequences [66]. The study presented a series of BERT models with varying parameter sizes. After pre-training on up to 250 million protein sequences, where each amino acid residue in a sequence was treated as a word, models accurately predicted the structural characteristics of proteins, including remote homology, secondary structure, and residue-residue contact. Representations emitted from the pre-trained 34-layer model were merged with multiple sequence alignments, which were the original input of existing secondary structure or contact prediction methods, and their prediction accuracy was improved. This result indicated that embedded representations from the pre-trained BERT incorporated more information than the multiple sequence alignments. Furthermore, the 34-layer model was fine-tuned to predict the quantitative effect of mutations and outperformed the state-of-the-art methods. As an attractive topic, other protein BERT models, such as TAPE transformer and ProtBert, have also been developed [65, 67]. Meticulous inspection of the TAPE transformer revealed that attention maps extracted from the pre-trained model reflect the context of input amino acid sequences [77]. For instance, one attention module, which specializes in deciphering residue-residue interactions, exhibited a significant correlation with experimental labels even though no structural information was provided. This phenomenon was later investigated by reconstructing protein contact maps from the attention maps of pre-trained ESM-1b [78]. The collection of studies illustrates that BERT-based models are highly interpretable and widely applicable to protein-related bioinformatics problems.

**DNABERT**, in contrast, is the only model, currently, to pre-train BERT-based models using a whole human reference genome [68]. During preprocessing, the genome, whose gaps and unannotated regions were excluded, was split into 5 to 510 consequent nucleotide sequences without overlapping and subsequently converted to 3-to 6-mer representations. Simply put, each subsequence of length 3 to 6 was regarded as a word. BERT models were pre-trained using k-mers with a masked language modeling objective and applied to downstream tasks. Upon task-specific finetuning, DNABERT demonstrated state-of-the-art or comparative performance in predicting promoter regions, binding sites of transcription factors (TFs), and splice sites. Attention analysis revealed that fine-tuned models captured the characteristics of each set of target sequences. For example, DNABERT fine-tuned using splicing datasets exhibited high attention weights in intronic regions in addition to target splice sites, indicating the ability of the model to learn the contextual significance of splicing enhancers or silencers in predicting splice sites. The study further applied DNABERT to predict promoters in the mouse genome and reported higher performance than those of existing deep learning methods. Overall, two-step training of the BERT architecture demonstrated its broad application to translate various genomic features in a cross-organism manner.

### 3.2. Applications for molecular interactions

Tsubaki *et al.* proposed a model by combining a graph neural network for compounds and a convolutional neural network (CNN) for proteins to predict compound-protein interactions (CPIs) [41]. Representations of compounds and proteins were obtained in an end-to-end manner. The word embeddings in the protein were learned from the training dataset using word2vec (3-mer of amino acids as words). To obtain protein vector representation, the average value of a set of hidden vectors was used with *d*-dimensional embedding after a hierarchical convolutional filter. Extensive evaluations were conducted on three CPI datasets (human, *C. elegans* [79] and DUD-E dataset [80]). The results showed that using the raw amino acid sequence as the input, the proposed approach significantly outperformed existing methods utilizing traditional chemical and biological features. They also established that the model could highlight 3D structural interaction sites between the compounds and proteins through an attention mechanism similar to that of words in sentences.

**ProbeRating** is a neural network-based recommender system utilizing word embeddings in NLP to infer binding profiles for unexplored nucleic acid-binding proteins (NBPs) [55]. ProbeRating achieves this goal using a two-stage framework. In the first stage, representation learning is performed using a package called FastBioseq, implementing FastText. Thus, the input feature vectors are extracted from the NBP sequences and nucleic acid probes. Authors chose 3-mers amino acids for proteins and 5-mers for nucleic acids as words. Three datasets (Uniprot400k [81], RRM3k [82], and Homeo8k [83]) were used to pre-train the FastBioseq protein embedding models, whereas RNA embedding models were trained directly from the RRM162 dataset [82]. In contrast, 8-mer frequency features were used for the DNA sequences in the Homeo215 dataset [84]. In the second stage, predicting the NBP binding preference was redefined as a recommender system formulation, where NBPs are like users and RNAs or DNAs are like products to be recommended. When no preference was available for a given user, the authors adapted and extended a strategy that converted the *binding intensity prediction* problem into a *similarity prediction* problem, solved it, and then converted it back. Extensive evaluation experiments were conducted on two tasks: RBP–RNA interaction and TF–DNA interaction. The results showed that ProbeRating outperformed three baseline methods (Nearest-Neighbor, Co-Evo [85] and AffinityRegression [84]). Further analysis suggested that this advantage was beneficial using both the neural network approach and input features extracted via word embeddings.

### 3.3. Applications in synthetic biology

Valeri *et al.* proposed a model that predicts synthetic riboregulators called toehold switches [86]. The model comprised a language model for toehold switch classification and a CNN-based model for toehold switch performance regression. In the language model, a sequence of toehold switches was embedded using ULMfit regarding a nucleotide as a word. They trained the model using toehold switches experimentally characterized by Angenent-Mari *et al.* [87]. The results showed that the model exhibited good and robust performance even for sparse training data and that the features obtained by the model revealed unknown properties of the toehold switches. They also showed that the trained model is easily fine-tuned by transfer learning using small external data [88, 89], and the fine-tuned model exhibited superior performance compared to an existing model. Finally, they showed that the fine-tuned model could help in the efficient design of toehold switches for various applications, such as SARS-CoV2 detection.

**UniRep** is a representation that comprehensively summarizes the semantics of arbitrary proteins and can be useful for various types of prediction tasks [90]. A protein sequence is embedded into UniRep using multiplicative LSTM (mLSTM), trained with 24M UniRef50 sequences [91], where an amino acid is regarded as a word. UniRep recapitulates biophysical properties, phylogenetics, and secondary structures of proteins. The authors also showed that UniRep outperformed other representations for predicting the structural and functional properties of *de novo* proteins, single point mutants, and natural proteins. These results suggest that UniRep is useful for the rational design of proteins. As a proof-of-concept, UniRep re-trained using deep mutational scanning data of GFP [92] was shown to effectively extrapolate GFP brightness outside the training domain. Therefore, UniRep was suggested to drastically reduce the cost for the rational design of GFP. Collectively, UniRep embodies various known protein characteristics and may be a versatile representation for protein bioinformatics.

### 3.4. Applications for other tasks

**StarSpace** is a *supervised* embedding method, which is different from the unsupervised embedding methods that we have introduced in section 2 [28]. Although StarSpace was originally developed for general NLP tasks, such as text classification, there are currently two bioinformatics applications. The first application is **BindSpace**, which predicts the binding sites of TFs [60]. BindSpace uses HT-SELEX experiments as the training dataset and applies StarSpace to the dataset by considering 8-mers and TFs as words and labels, respectively. In performance evaluation using the ENCODE ChIP-seq dataset, BindSpace achieved high classification performance even between paralogous TFs, which have highly similar binding motifs. The second application is **MutSpace**, which estimates the cancer types of patients from somatic mutation patterns [61]. This method regarded mutation patterns and cancer types as words and labels, respectively. MutSpace shows state-of-the-art performance in a breast cancer subclass classification problem. The high performance of these two applications means that StarSpace is likely to perform well in other bioinformatics problems.

A constrained semantic change search (**CSCS**) is a method for discovering word changes that significantly alter the semantics from an original sentence based on embedding techniques [93]. The key feature of this method is that it does not detect word changes that would abolish the grammar of the sentence but those that preserve the grammatical structure. For example, in an NLP task, CSCS can change “winegrowers revel in *good* season” to “winegrowers revel in *flu* season.” We briefly introduce the CSCS method. We define *x* and 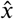 as the original and mutated sentences, respectively. The embedded representations of *x* and 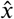 are defined as *z* and 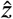, respectively. Here, the semantic change is modeled as the distance between these embedded representations, that is, 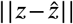. Additionally, the preservation of the grammatical structure is evaluated by 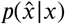, which is also modeled using embedding techniques. Finally, 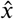 maximizing 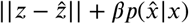, where *β* is a scaling factor. One biological application of CSCS is the modeling of viral evolution [56]. This application regarded viral proteins, preservation of the infectivity, and escape from antibody recognition as sentences, preservation of grammar, and semantic change, respectively, and detected escape mutations from immune systems as a result of the CSCS analysis. The analyses of HIV-1 and influenza viruses showed that mutations detected by the CSCS were in good agreement with the experimental mutation results.

Woloszynek *et al.* applied word2vec to a metagenomic dataset by regarding *4–15-mers* in sequencing reads as words [38]. They trained word2vec with a skip-gram model using 2,262,986 full-length 16S rRNA amplicon sequences from GreenGenes [94], a microbial 16S rRNA sequence database obtained using metagenomic analysis. They verified the robustness of the model in a taxonomic identification task using an independent dataset of 16,699 full-length 16S rRNA sequences from the KEGG REST server [95] as a validation dataset. The embedding features exhibited superior performance to the k-mer frequency features. In addition, the embedding as also performed using the American Gut project dataset [96], which has 11,341 partial 16S rRNA sequences from three body sites (gut, skin, and oral cavity), and showed comparable performance to conventional methods, such as sequence alignment in the body site classification task. These results suggest the availability of embedding with pre-trained models instead of sequence alignment for metagenomic sequence profiling.

## 4. Summary and outlook

In this study, we introduced basic algorithms and reviewed the recent literature concerning representation learning applications in sequence analysis. Heinzinger, *et al.* pointed out three difficulties in biological sequence modeling with NLP [62]as follows: (i) Proteins range from approximately 30 to 33,000 residues, which is much longer than the average English sentence, which consists of 15 to 30 words [97]; (ii) proteins use only 20 amino acids in most cases; if we consider one amino acid as a word, the word repertoire is 1/100,000 of English language, and if we consider 3-mer as a word, the word repertoire is 1/10 to 1/100 of English language; (iii) UniProt is more than ten times the size of Wikipedia, and extracting information from a huge biological database may require a commensurate model. Embedding biological sequences using NLP overcomes these difficulties and outperforms existing methods in many tasks, such as function, structure, localization, and disorder prediction (Table 1). In addition to these general biological tasks, representation learning has also been used to solve specific problems, such as RNA aptamer optimization [98], viral mutation prediction [56], and venom toxin prediction [36]. In these studies, representation learning of biological sequences could capture biophysical and biochemical properties of biological systems, and representation learning may reveal the grammar of life.

The development of novel representation learning methods has been actively studied in machine learning research. For example, hyperbolic embedding methods have been pursued in recent years [99, 100]. These methods embed the data not in Euclidean space, which is utilized in all the studies introduced in this paper, but in the *hyperbolic* space. The hyperbolic space has constant negative curvature; thus it shows characteristic geometric features not seen in Euclidean space, such as the sum of the interior angles of a triangle being less than 180°. Changes in the embedding space can considerably alter the efficiency of representation learning, and theoretical and experimental analyses have shown that hyperbolic embedding methods are suitable for data with hierarchical latent structure. Therefore, hyperbolic embedding methods have recently been used for biological analysis, such as phylogenetic analyses [101] and single-cell RNA-seq analyses [102]. Furthermore, research on embedding into more complex spaces, such as mixed-curvature spaces, has also attracted attention [103]. The application of these embedding techniques in non-Euclidean space for biological sequence analyses should be an essential research direction in the future.

New approaches are released every day in this field, and the scientific community is trying to compare their accuracy and validate their uses [65,104,105]. Therefore, it is important to make the models available in an easy-to-use form with documentation. In addition, considering the rapid growth of biological databases, the source code for creating models should be made available for future updates. Only a limited number of studies have released both the source code and the pre-trained model with the relevant documentation. Participants in this community need to publish their papers in a form that can be reproduced and verified.

In this study, we comprehensively surveyed and reviewed the application of representation learning to biological sequence analysis. Although NLP-based biological sequence analysis is still in its early stages and requires further development, in the light of novel challenges in biology, such as single-cell analysis, genome design, and personalized medicine, representation learning may help the progress of bioinformatics studies.

## Acknowledgements

The illustrations in Figure 1 were kindly provided by Kae Namie. This work was supported by the Ministry of Education, Culture, Sports, Science, and Technology (KAKENHI) [grant numbers: JP17K20032, JP16H05879, JP16H06279 JP19H01152 and JP20H00624 to MH, JP19K20395 to TF, JP19J20117 to SH and JP20J20016 to TM] and JST CREST [grant numbers: JPMJCR1881 and JPMJCR21F1 to MH].

